# Behind the shadow play: Shedding light on the population genetics of partially clonal organisms through clone age distributions

**DOI:** 10.1101/2025.10.03.680262

**Authors:** Ammar Abdalrahem, Camille Noûs, Sébastien Duplessis, Pascal Frey, Solenn Stoeckel, Fabien Halkett

**Author notes:** Corresponding author: Fabien Halkett, Ammar Adbalrahem. These authors contributed equally to this work (Solenn Stoeckel, Fabien Halkett).

## Abstract

Partial clonality is widespread in natural populations; however, the distributions of clone ages under different rates of clonality and their effects on genetic diversity remain unexplored. To fill this gap, we simulated partially clonal populations over 10,000 generations across a range of clonality and mutation rates. Using a forward-in-time individual-based model, we evaluated genetic and genotypic indices alongside measures of clone age distribution to examine how different clonality rates influence clone age distributions, and how these distributions impact population structure and genetic diversity over time.

Our results reveal two distinct trajectories: (i) at low to moderate rates of clonality evolution is driven by sharp and predictable patterns of rapid clone turnovers, resulting in predominantly young clones which relative abundances well align with genotypic indices ; (ii) at extreme rates of clonality, demographic stochasticity generates very variable clone age distributions which mirror the high variance of mean and variance of *F*_IS_ that summarize gene reshuffling between individuals. Interestingly, a portion of the variability of these indices can be explained by differences in clone age distribution that occur by chance. Our results, therefore, complement previous theoretical studies on the population genetics of partial clonality by providing biological insights into how clonal turnover dynamics shape the temporal evolution and variability of highly clonal populations. These results have practical implications for inferring evolutionary trajectories and managing partially clonal species.

## Introduction

Clonal reproduction is common throughout the Tree of Life. It has profound consequences for the evolution of organisms and shapes genetic diversity within and between populations (Rozenfeld et al., 2007; Tibayrenc et al., 2015; Stoeckel et al., 2021). In particular, it distorts genetic equilibrium through the burst of clones (Arnaud-Haond et al., 2007). However, how clonal turnover (that is, the distribution of clone ages) structures genetic diversity remains largely unknown.

A diverse array of reproductive strategies exists, ranging from obligate sexuality to strict clonality, with many species employing mixed reproductive modes, which is often referred to as facultative sexual reproduction or partial clonality (Halkett et al., 2005; Hartfield, 2016; Stoeckel, et al., 2021). In the following, we use the latter expression, as it has broader acceptance in both experimental and theoretical literature. Partial clonality is a common reproductive strategy whereby individuals are capable of both sexual reproduction and clonal propagation. In these populations, genetic individuals (genets) persist across generations through asexual ramet production, while sexual recruitment introduces novel genets via recombination (Hartfield, 2016; Stoeckel, Porro, et al., 2021; Orive & Krueger-Hadfield, 2021). Clonality (also sometimes named asexuality) refers to the reproduction of identical individuals (with the exception of somatic mutations) from one generation to the next. Orive & Krueger-Hadfield (2021) detail a lexicon of species-specific means to achieve clonal reproduction. Partial clonality has been documented in many plant species, such as seagrasses and duckweed *Spirodela polyrhiza* (Ho et al., 2019; Arnaud-Haond et al., 2020), invertebrates like aphids and coral reefs (Halkett et al., 2005; Carpenter et al., 2008), fungi like cereal rusts (Pucciniales) (Ali et al., 2014; Duplessis et al., 2021) and pathogenic yeasts (e.g. *Candida albicans*) (Nébavi et al., 2006), and protozoan parasites such as *Plasmodium falciparum* (Early et al., 2022). These organisms exhibit a diverse range of ecological niches, spanning from primary producers to the highest trophic level, and a large array of lifestyles, including free-living organisms but also many plant or animal (including human) pathogens.

Partial clonality impacts the ecological and evolutionary trajectories of species populations. On one hand, clonal reproduction leads to rapid demographic expansion, exploiting stable environments, without paying the twofold cost of (biparental) sex (Lehtonen et al., 2012; Lodé, 2013). On the other hand, sexual reproduction results in allelic shuffling through recombination, which is crucial for adaptation (Roze, 2012; Pierre et al., 2022). Partial clonality could take advantage of the benefits of each reproduction mode, as evidenced for many parasites and pests (McDonald & Linde, 2002; Bazin et al., 2014). For instance, *Phytophthora infestans*, the oomycete responsible for the Irish Potato Famine by causing potato late blight disease, relies on clonal reproduction to spread rapidly, while sexually reproducing to generate new lineages, posing an ongoing threat to global agriculture (Gavino et al., 2000; Fry et al., 2015). Malaria is a human life-threatening disease caused by the *P. falciparum*. The infection is dominated by rapid clonal multiplication within human red blood cells, while sexual reproduction occurs in the mosquito vector (Chang et al., 2013; Early et al., 2022).

Genotypic and genetic measures are crucial for understanding the evolutionary trajectory and ecological potential of organisms that reproduce both clonally and sexually (Stoeckel, Arnaud-Haond, et al., 2021; Stoeckel, Porro, et al., 2021). Clonal lineages can serve as an “evolutionary memory” of the population’s history, maintaining genetic signatures of past events, such as bottlenecks and founder effects, through the freezing of ancestral genotypes (Brakefield, 1989; Reichel et al., 2016). Theoretical and empirical studies indicate the existence of clonality within the population through genotypic metrics using traditional and genome-wide molecular markers (e.g., SSR and SNPs, respectively) (Stoeckel, Porro, et al., 2021; Abdalrahem et al., 2025), such as clonal richness (*R*), which estimates the proportion of unique multilocus genotypes (MLGs), and the Pareto β index, which estimates the distribution of clonal sizes, highlighting the dominance of certain MLGs (Arnaud-Haond et al., 2007). Furthermore, genetic measures such as the inbreeding coefficient (*F*_IS_), where negative values may indicate an excess of observed heterozygotes (*H*_O_). Clonality often increases linkage disequilibrium (LD), indicating the presence of allelic associations due to limited recombination among lineages (Balloux et al., 2003; Halkett et al., 2005; Stoeckel & Masson, 2014).

Clonality within partially clonal populations slows down the convergence to the Hardy-Weinberg equilibrium (HWE) state, creating temporary deviations in genetic patterns that influence both genotypic and genetic indices (Reichel et al., 2016). This “transient” genetic deviation highlights the importance of estimating the rate of clonality (*c*) to measure the proportion of clonal and sexual reproduction within populations, which influences our interpretation of genetic diversity and equilibrium dynamics (Stoeckel, Porro, et al., 2021). For example, at low *c*, populations often follow HWE expectations, and the reduction of genotypic diversity indicates historical bottlenecks or founder effects (Hanko et al., 2023), but at high *c*, it may indicate dominance by a few clonal lineages (Reynes et al., 2021; Abdalrahem et al., 2025). Moreover, a negative *F*_IS_ value is the hallmark of a high clonality rate (Balloux et al., 2003; Stoeckel & Masson, 2014), although slightly negative *F*_IS_ value can also be observed for outcrossing species (Cabrera-Toledo et al., 2008). Although the effect of clonality rate has been largely established theoretically (Balloux et al., 2003; Halkett et al., 2005; Stoeckel & Masson, 2014; Becheler et al., 2017; Stoeckel, Porro, et al., 2021), it is seldom used in empirical studies dealing with partially clonal organisms, leading to potential misinterpretations of key population genetic indices. Here, we propose to better highlight the link between the rate of clonality (*c*) and the variation of population genetic indices by examining the clonal turnover, which denotes the pace at which alleles are reshuffled within and between individuals.

Despite the recognition of the influence of partial clonality on population genetic structure, the evolutionary consequences of clonal age distributions (i.e., the variation and structure of clone longevity) remain unexamined across rates of clonality, a critical gap given its implications for mutation accumulation and adaptive potential. Early work proposed that clone age, due to mutation accumulation, would profoundly impact genetic diversity (the ’Meselson effect’; Birky, 1996; Mark Welch & Meselson, 2000; Balloux et al., 2003), yet no study addresses how increasing *c* values reshape clone age distributions or their downstream diversity effects. We hypothesize that high rates of clonality shift age structure toward older lineages, amplifying mutation accumulation and decoupling genotypic from allelic diversity. Specifically, increased *c* values may reduce genotypic richness (*R*) through dominance of persistent clonal lineages while paradoxically maintaining allelic diversity (*H*_E_) through genotype freezing that prevents genetic drift.

To resolve this gap and test our hypothesis, we employ individual-based forward simulations for a first exploration on how clonality rate (*c*), mutation rate (*μ*), and population size (*N*) jointly shape genetic indices and clone age distributions over generations. Our aim here is to disentangle the deterministic effects of clone age distribution on population genetic indices from its random component due to the stochasticity of their evolutionary trajectories using 100 simulation replicates. We simulate populations under an annual life cycle where each discrete generation corresponds to one reproductive season. Our model tracks clone age distributions across the full range of rates of clonality, from strictly sexual (*c*=0) to strictly clonal (*c*=1) reproduction, over 10,000 generations, while dynamically quantifying genetic diversity indices and genotypic metrics at regular time points. In the following, we first revisit the effect of clonality rate (c) on the temporal variation of population genetic indices, focusing on two iconic mutation rates (classically set at the scale of the genetic drift i.e. *μ=*1/N, corresponding here *μ=*10^-3^ for a population size N=1000 or at a reduced but biologically relevant mutation rate of 10^-6^ thus when genetic drift is dominant). Then, we characterize the changes in clone age distributions according to clonality rate (*c*) and the variance inflation in clone age structure, looking if high variances of genetic indices that correspond to some diffuse distributions of clone age. Last, we examine the correlations between descriptors of clone age distributions and the values of population genetic indices obtained both within and among *c* values. The strengths and shapes of these correlations are discussed from the perspective of inferring clone turnover in empirical studies.

## Materials and Methods

### Simulation design

We performed individual-based, forward-in-time simulations to investigate how clonality rate (*c*: proportion of clonal reproduction) and mutation rate (*μ*: mutations per locus per generation) shape population genetic indices under clone age structure dynamics (Fig. 1). Each simulation modeled a population of diploid individuals with a constant size (N = 1,000 individuals) maintained across generations, thereby isolating the effects of *c* and *μ* on genetic diversity, while avoiding confounding demographic fluctuations (Gazave et al., 2013; Raynes et al., 2018). Reproduction occurred via clonal propagation (producing genetically identical offspring) and/or sexual outcrossing (generating recombinant offspring), with the relative contribution of each mode controlled by *c*.

**Fig. 1.**
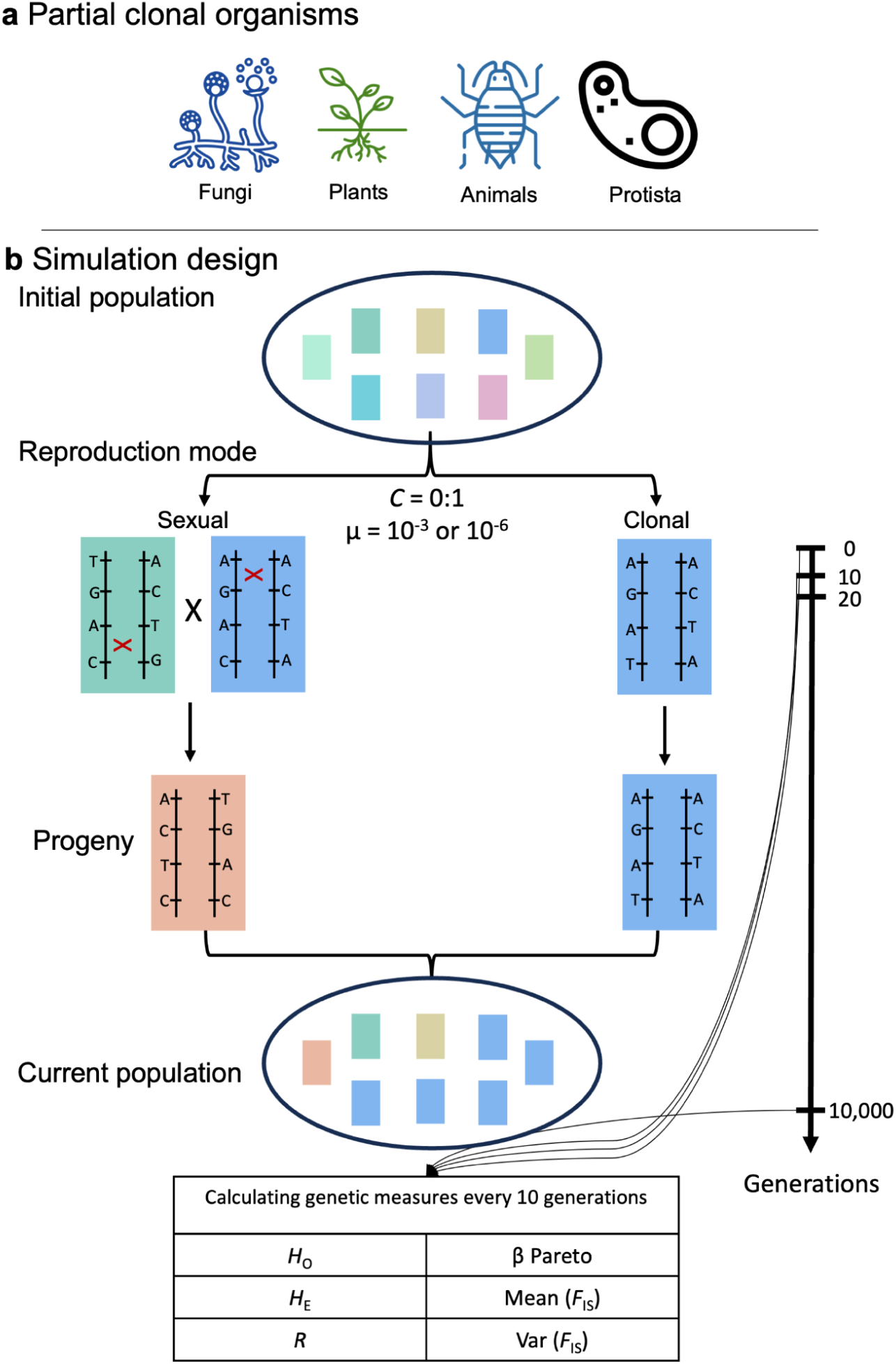
Simulation workflow for investigating clone age structure and genetic diversity in a partial clonal system. The simulation begins with a population of 1,000 diploid individuals at the Hardy-Weinberg equilibrium (HWE). Each generation, individuals reproduce either sexually or clonally, depending on the clonality rate (c). Mutation is introduced per locus at rates of μ = 10⁻^3^ or μ = 10⁻⁶, with each locus initially harboring up to four alleles. The simulation proceeds for 10,000 generations with 100 replicate runs per parameter combination. Population genetic indices (e.g., inbreeding coefficient (F_IS_), expected heterozygosity (H_E_), mean observed heterozygosity (H_O_), clonal richness (R), Pareto β index), calculated every 10 generations.

We explored rates of clonality ranging over 19 fixed values, spanning from fully sexual (*c* = 0) to fully clonal (*c* = 1), with increased resolution at high *c* values (0.95, 0.96, …, 0.9999) to capture thresholds where clonal dominance alters genetic dynamics. We implemented a k-allele mutation model (KAM) (Bhargava & Fuentes, 2010; Putman & Carbone, 2014), where each locus can harbor up to a set number of distinct alleles per locus (Nballele), and mutations transition stochastically between these alleles at rate *μ* (Stoeckel, Porro, et al., 2021). For that, we varied the number of alleles (Nballele) to 2 (biallelic), 4, 8, or 99 (approximating an infinite alleles model) to capture differing levels of allelic richness. To ground simulations in realistic systems, we focussed our analyses on two mutation rates: μ = 10⁻⁶ representing the mutation rate of a genome-wide marker data, such as single-nucleotide polymorphisms (SNPs), and μ = 10⁻³ as the rate of a traditional marker data like microsatellites (SSR) (Payseur & Cutter, 2006; McCulloch & Kunkel, 2008; Stoeckel, Porro, et al., 2021). All loci evolve independently, exhibiting no linkage, and are selectively neutral to sort out the effects of c and μ on population genetic indices.

Simulation ran for 10,000 generations, with genetic data collected at 10-generation intervals (0, 10, 20, …, 10,000) to resolve transient and equilibrium states and reduce data redundancy. For each parameter combination (c, μ, Nballele), 100 replicate runs were performed to quantify stochastic variance, which is sufficient to capture 95% confidence intervals for diversity metrics (Rijnsoever, 2017). Simulation was implemented in a custom Python 3 script adapted from Stoeckel et al. (2021), which tracked clone ages (the number of generations a clone has maintained clonally in the population, corresponding to the number of generations since this genotype was produced by a sexual event) and evolutionary parameters including μ, c, and generational transitions (Fig. 1).

### Analysis of population genetic indices across rates of clonality

As outputs, we calculate several population genetic indices for each replicate. The mean expected heterozygosity (*H*_E_) and mean observed heterozygosity (*H*_O_) were calculated as measures of genetic variation at the allele level (Nei, 1978). Additionally, the mean and variance of the inbreeding coefficient (*F*_IS_) were computed, indicating the deviations from the Hardy–Weinberg equilibrium (Wright, 1931, 1949; Stoeckel & Masson, 2014). The total number of alleles and the number of fixed loci were also captured as summaries of allelic diversity and fixation dynamics. To assess clonal structure, the total number of distinct genotypes, referred to as clonal richness (*R*), was derived as an index of clonal diversity, and Pareto *β* was estimated to capture the distribution of genotype frequencies (Arnaud-Haond et al., 2007). We used the Python library from GenAPoPop software (Stoeckel et al., 2024) to compute population genetic indices from the simulated data. Clone age structure was captured by recording the number of individuals of each age (clone age abundance) for each simulation replicate. By definition in our system, the age of all individuals produced by a sexual event in the last considered generation was set to zero.

### Temporal analysis of genetic indices across clonality rates

To analyze the temporal dynamics of population genetic indices under different rates of clonality, we filtered the simulation data to focus on a representative set of *c* values: 0.1, 0.3, 0.5, 0.7, 0.9, 0.95, 0.98, 0.99, 999, and 0.9999. This selection aims to improve clarity in visual comparisons of the key transitions of the dynamics of reproductive modes within populations and their impact on genetic structure. We summarized the trend in the evolution of population genetic indices (calculated using the methods described above) by calculating the median value across all simulation replicates for each combination of clonality rate, mutation rates (10^-4^ or 10^-6^), at each time point (generation).

After quantifying the temporal dynamics of population genetic indices across generations, we selected one generation (time point) to examine the relationship between population genetic indices and the distribution of clone ages. We set this generation in such a way as to tip the balance towards describing a stable distribution for most population genetic indices, while limiting the effect of allele fixation over time.

### The effect of rates of clonality on clone age distribution

For each replicate, we recorded the number of individuals associated with each clone age at the end of the simulation (I.e., after 10,000 generations). We analysed clone age and abundance of clone ages with varying rates of clonality and mutation rates to assess how the persistence and prevalence of clones of different ages varied. Overall distributions of clone ages for each rate were visualised using joy plots (ridge plots) to compare age structure patterns.

The distribution of clone ages for each replicate was then summarized using basic descriptors :

- the median, maximum, and variance of clone age distribution.

- the slope and the coefficient of determination (R²) values of the linear regression between the logarithm of the abundance (number of individuals) and clone age. These last two descriptors encapsulate both the shape (slope) and the predictability (R²) of the relationship between clone age and abundance across different rates of clonality.

To examine how clone age-abundance relationships relate to population genetic structure, we computed a pairwise Spearman correlation coefficient matrix using pairwise complete observations. At the fixed generation, we merged the regression outcomes (slope and R²) values with corresponding genotypic and genetic indices for each simulation replicate, including the mean and variance of inbreeding coefficients (*F*_IS_), observed and expected heterozygosity (*H*_O_, *H*_E_), clonal richness (*R*), and the *Pareto β* parameter. This indicates the strength and direction of monotonic associations between the regression outcomes and the various population genetic indices. Additionally, we generated scatter plots to visually explore the relationships between each clone age-abundance metric (slope and R²) and the population indices. These plots enabled us to detect potential nonlinear patterns or outliers that may not be apparent from correlation coefficients alone.

## Results

### Introducing the effect of rates of clonality on the temporal change of population genetic indices

First, we analyzed the dynamics of variation of population genetic indices over generations (time). By tracking these indices across 10,000 generations, we shed light on how their values dynamically change in response to both clonality rate (*c*) and mutation rate (*μ*). This approach captures the overall patterns of temporal changes in both genotype composition and allelic states within the population, providing a comprehensive view of how partial clonality shapes population genetics across diverse evolutionary scenarios (Fig. 2).

**Fig. 2.**
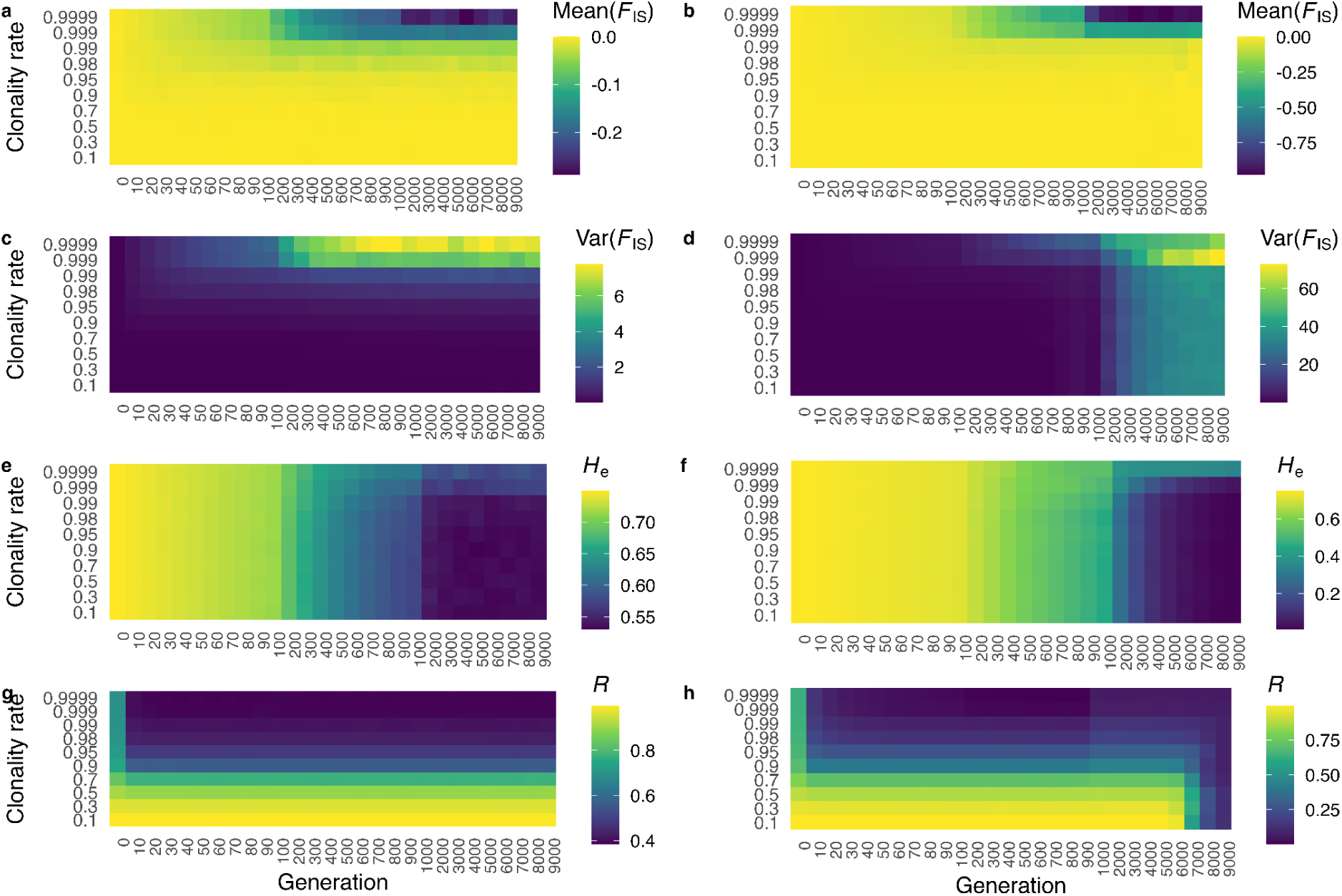
Effect of clonality and mutation rates on genetic and genotypic diversity. Heatmaps showing the effects of clonality rate (*c*) and mutation rate (*µ*) on genetic and genotypic diversity indices across simulated generations. The left column (a, c, e, g) presents results for a high mutation rate (μ = 10⁻^3^), while the right column (panels b, d, f, h) represents a low mutation rate (μ = 10⁻^6^). From top to bottom, the panels represent: mean value of inbreeding coefficient (Mean (*F*_IS_)), variance in inbreeding coefficient (Var (*F*_IS_)), expected heterozygosity (*H*_E_), and clonal richness (*R*). Within each graph, the y-axis represents the rate of clonality (*c*), ranging from 0.1 to 0.9999, and the x-axis shows the generation number (0 to 9,000). Color intensity indicates the mean value over 100 simulation replicates, with yellow representing higher and purple lower values. Note the difference in scale values between mutation rates.

At low to moderately high clonal rates (*c* ≤ 0.95), the mean inbreeding coefficient Mean (*F*_IS_) at a mutation rate of μ = 10⁻^3^ (Fig. 3a) remains close to zero or slightly positive, reflecting minimal deviation from HW expectations over generations even when populations are observed for up to 9,000 generations. However, at higher clonality rates (c ≥ 0.98), a pronounced temporal shift emerges, with Mean (*F*_IS_) becoming strongly negative after generation 1,000. When the mutation rate is low (μ = 10⁻^6^), hence genetic drift is the dominant force, a similar qualitative trend is observed, but Mean (*F*_IS_) becomes even more negative at very high clonality rates (c ≥ 0.999). Then, the scale of mean (*F*_IS_) differs between mutation rates, reaching values close to -0.8 at low mutation rates and around -0.3 at high mutation rates. This illustrates that mutation accumulation is not the main driver of heterozygote excess in partially clonal populations. Values of mean (*F*_IS_) have to be examined in the light of the marked genetic drift that applies at a low mutation rate (μ = 10⁻^6^). Except for very high rates of clonality, gene diversity (*H*_E_) collapses. After generation 6,000, most loci tend to become fixed, and genetic diversity is only preserved within the genotypes frozen by clonal reproduction, when loci are heterozygous by chance, which generate these extreme values of mean (*F*_IS_), almost artefactually.

**Fig. 3.**
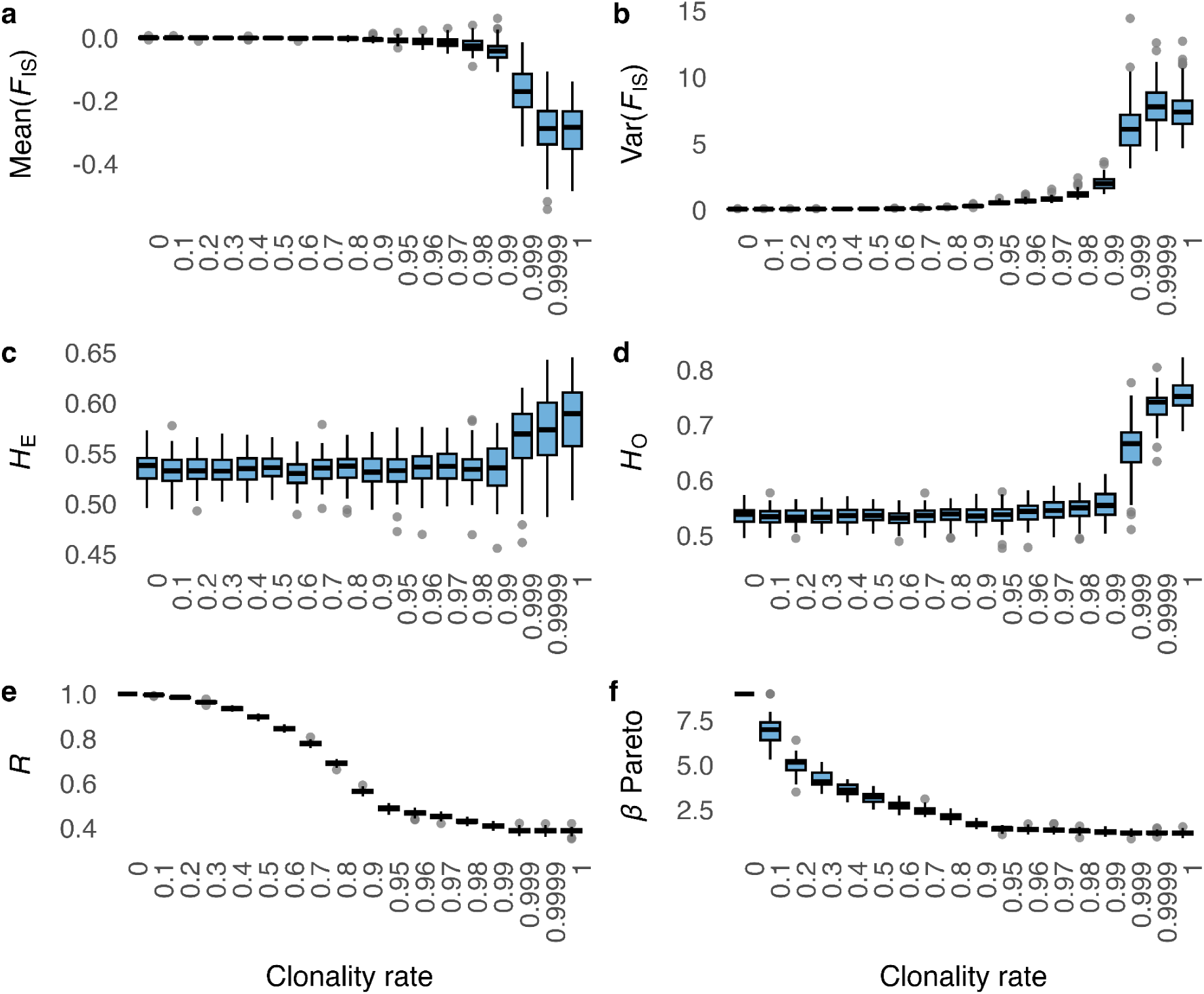
Effects of clonality rate on population genetic indices. Boxplots show the distribution of population genetic indices based on 100 simulation data replicates per different clonal rates *c*, at generation 6,000. The population genetic characteristics: (a) mean, and (b) variance of the inbreeding coefficient *F*_IS_, (c) mean expected heterozygosity *H*_E_, (d) mean observed heterozygosity *H*_O_, (e) clonal richness *R*, (f) Pareto *β* index The simulation was conducted with a population size of N=1,000, 100 loci, mutation rate *μ=* 10^-3^, and four alleles.

Notably, the variance of the inbreeding coefficient, Var (*F*_IS_), exhibits a slightly different pattern, with a difference between mutation rates. Under a high mutation rate (μ = 10⁻^3^, Fig. 3c), the variance remains relatively low across generations, rising moderately to values between approximately 2 and 6 at extreme clonality (*c* ≥ 0.999). In contrast, under a low mutation rate (μ = 10⁻^6^, Fig. 3d), Var (*F*_IS_) stays consistently low over most clonality rates and generations, but exhibits a notable increase after about 1,000 generations. At these later stages and higher clonality, Var (*F*_IS_) escalates substantially, reaching much larger values ranging roughly from 20 to 60. In accordance with previous theoretical studies, Var (*F*_IS_) does not increase monotonously with clonality rate, the highest value being observed for very high (*c* = 0.999) but not the most extreme c values (*c* ≥ 0.9999). Clonal richness (*R*) measured over the entire population exhibited distinct temporal patterns that varied with both clonality rate and mutation rate (Fig. 3g, h). Initially, all populations began with high *R* values across the clonality gradient, but these *R* values, computed on all individuals of a population, almost immediately aligned with the rates of clonality. For moderate to high clonality rates (c ≥ 0.5), *R* values declined sharply within the first 10 generations under both mutation rates, decreasing progressively with increasing c, reflecting that the population is composed of fewer clonal lineages. At a high mutation rate (μ = 10⁻^3^, Fig. 3g), this pattern is invariant throughout the 9,000 generations. At low mutation rate (μ = 10⁻⁶, Fig. 3h), low to moderate clonality rates (*c* = 0.1: 0.7) maintained relatively high clonal richness (R ≈ 0.75) up to generation 5,000, but declined after generation 6,000, reaching R ≈ 0.25 by generation 9,000. Here, again, this is an effect of genetic drift that causes a loss of discrimination among individuals because of a reduced set of polymorphic loci. This difference in scale of population indices values reflects how the mutation rate interacts with clonal reproduction to shape the genetic structure and variability over time.

In the following, we focus our analyses on the highest mutation rate (μ = 10⁻^3^), since artifactual variations of population genetic indices are observed for a low mutation rate (μ = 10⁻⁶). We chose to consider generation 6,000 as a good compromise between observing very old clones while not enabling genetic drift to fix only one clone at high rates of clonality.

### How rates of clonality drive the variability in evolutionary trajectories

To document the effect of increased rates of clonality on the stochastic variation in population genetic indices, we analyzed the genetic states of populations observed at a single time point (generation 6,000), considering a mutation rate of μ = 10⁻³ and four alleles per locus (Fig.3). Notably, the variability among replicates of both the mean and variance of inbreeding coefficient (*F*_IS_) across loci increases at high rates of clonality (*c* > 0.99), when simulation departs from Hardy-Weinberg expectations. The variations of expected heterozygosity (*H*_E_) and observed heterozygosity (*H*_O_) remain relatively even, with only a slight increase at higher clonality rates (*c* ≤ 0.999), more pronounced for observed heterozygosity (*H*_O_). Notably, the variability of *H*_O_ is maximal at *c* = 0.999, with simulation replicates that either display values consistent with moderate or high clonality rates.

The clonal richness (*R*) and the clonal evenness (*Pareto β*) are both inversely related to the rate of clonality, but with marked differences at low to moderate *c* values. Whereas *R* displays a bell-shaped curve decreasing with *c* values, with no difference in variability across *c* values, *Pareto β* displays an exponential decrease with marked variability among replicates at low clonality rate (*c* ≤ 0.3). The variation in these indices indicates that the clonal structure varies greatly even at low values of *c*.

### The rate of clonality shapes clone age structure

To investigate the impact of the rate of clonality on population age structure, we examined the distribution of clone ages across a broad gradient of clonality rates throughout 10,000 generations (Fig. 4). At low to moderately high clonality (*c* ≤ 0.8), clonal age was limited, with clones rarely surviving more than 100 generations. Even at very high clonality (*c* = 0.99), maximum clone age remained below 1,000 generations. In contrast, at *c* = 0.999, a different pattern emerged, with a more flattened distribution of clone ages. Here, the population is composed of a mixture of clones ranging from quite young to much older, with some clones that could persist by chance for over 9,000 generations. When *c* > 0.999, the population is mostly composed of very old clones, with the extreme case of *c* = 1 in which the population consists solely of clones of the maximum age of 10,000 generations. This sharp transition across the gradient of clonality rates demonstrates that while increasing clonality gradually extends clone ages, the turnover of clones remains very high. Extremely high clonality is required to enable the long-term persistence of clones.

**Fig. 4.**
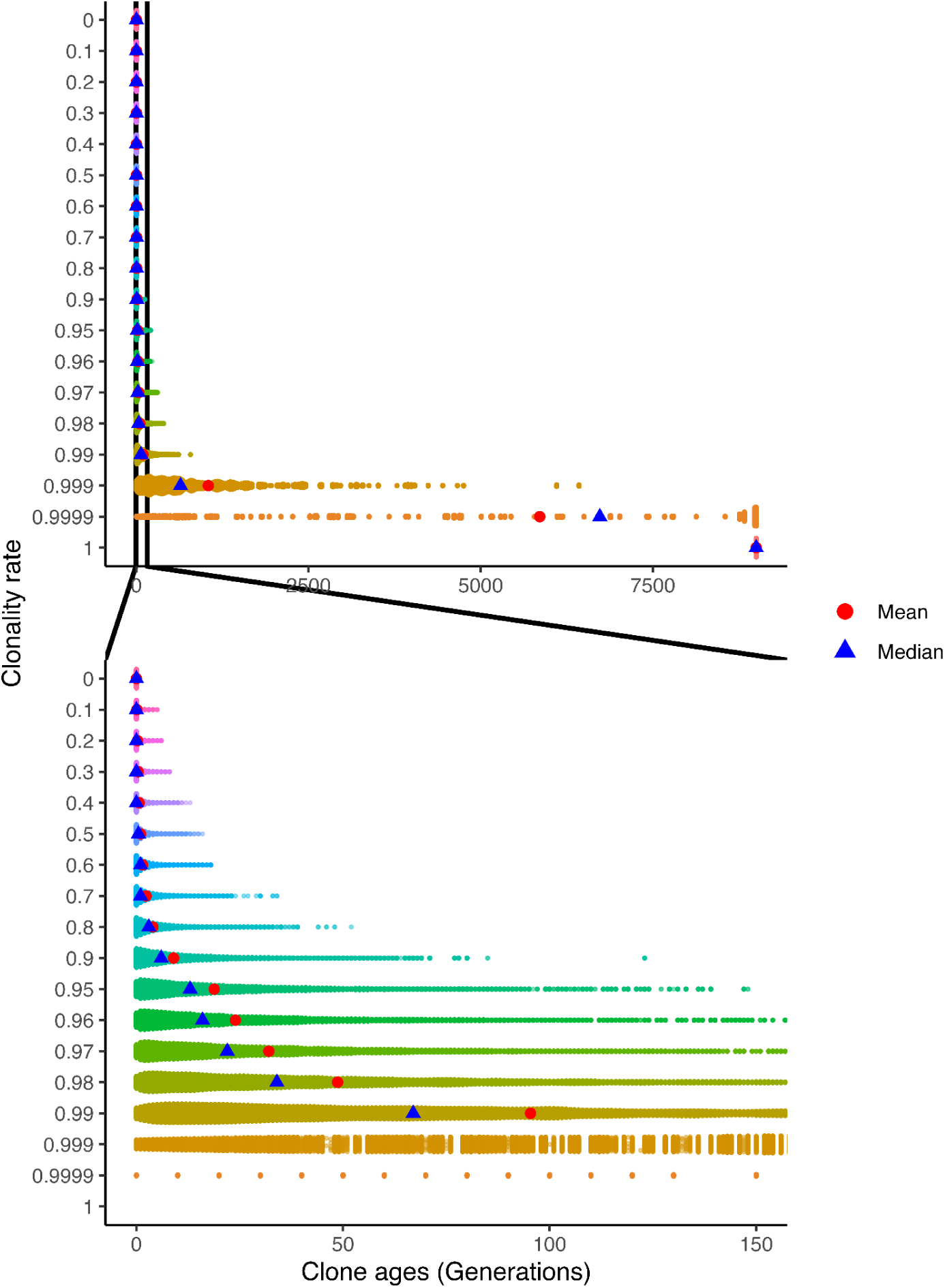
Clone age distributions across clonality rate gradients. Joyplot showing the distribution of clone ages (generations) across clonality rates. Each density curve represents the age distribution of clones at a specific clonality rate (*c*), with curve height indicating the distribution of individuals at each age. Below the main graph is the zoom-in on the first generations (0 to 150) for clearer visualization of the onset of the shift of mean and median clone age.

### The abundance of clones responds to changes in rates of clonality

To examine the relationship between clone age and clone abundance of individuals under a gradient of *c* values, we analyzed their joint distribution at generation 9,000 (Fig. 5). At low to moderate clonality rates (*c* ≤ 0.7), nearly all clones were young, with few individuals persisting beyond 30 generations. Importantly, these clones showed a visible pattern of the distribution of abundance according to their ages, with the logarithm of abundance that fits a linear regression with age (Fig. S1 & S2).

**Fig. 5.**
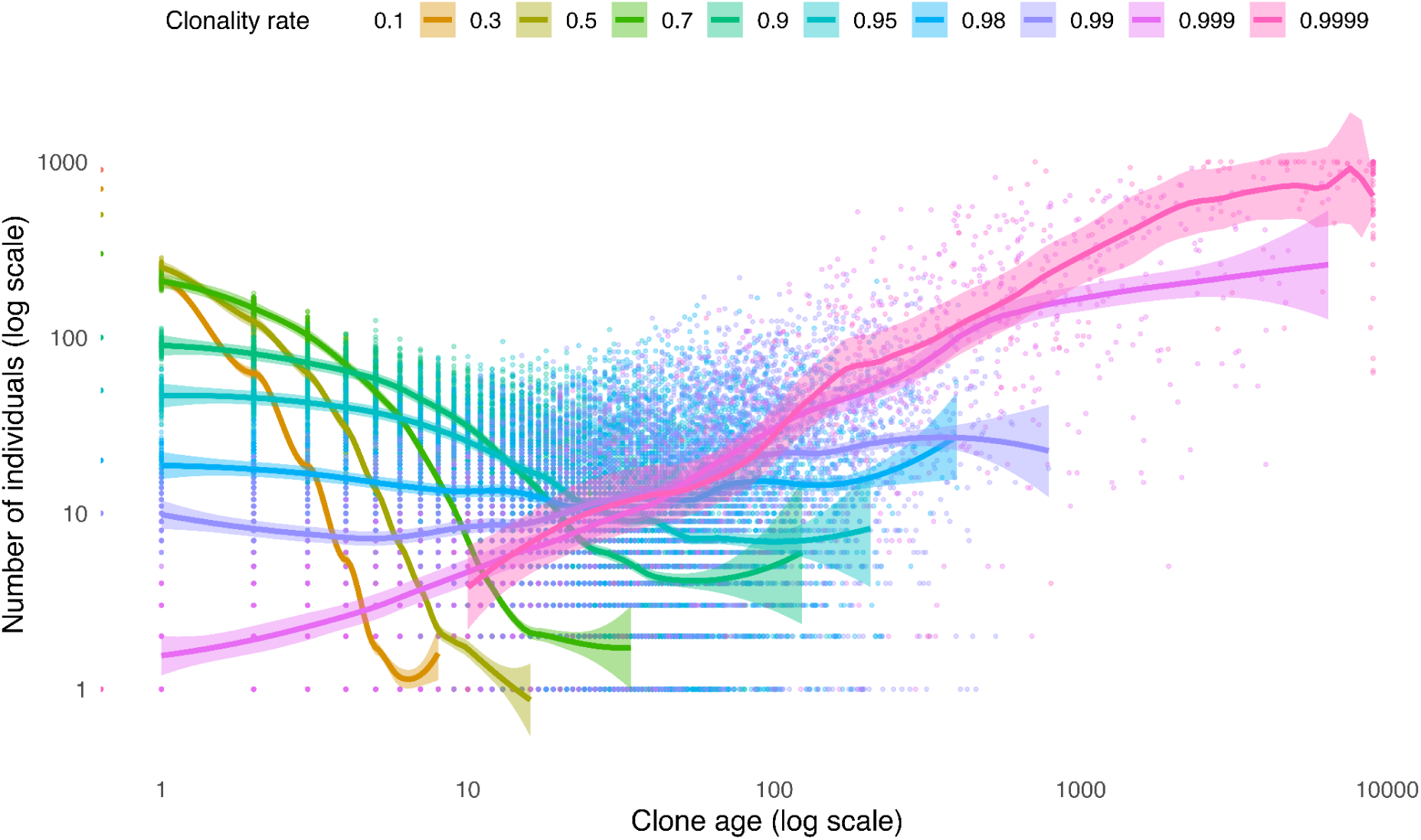
Clonal age distribution across a gradient of clonality rates. The plot shows the relationship between clone age (log scale, in generations) and clone size (log scale, number of individuals per clone) at generation 9,000. Each point represents a distinct clone observed in the population at this time. Colored LOESS curves (span = 0.3) illustrate smoothed trends for different clonality rates (*c*), with shaded ribbons indicating standard errors.

However, at higher clonal rates (*c* ≥ 0.98), two key shifts emerged: 1) populations contained numerous large clones persisting for hundreds of generations, and 2) the distribution became diffuse, with clone ages and sizes exhibiting a stochastic rather than a structured correlation pattern as observed at lower clonality rates (Fig.S3). This diffusion indicates a change in age-abundance distribution patterns at extreme clonality, where the structured relationships observed at lower clonality rates become unpredictable (Fig. 6c). Moreover, the greatest number of clone age classes was observed at the diffusion point (Fig.S5).

**Fig. 6.**
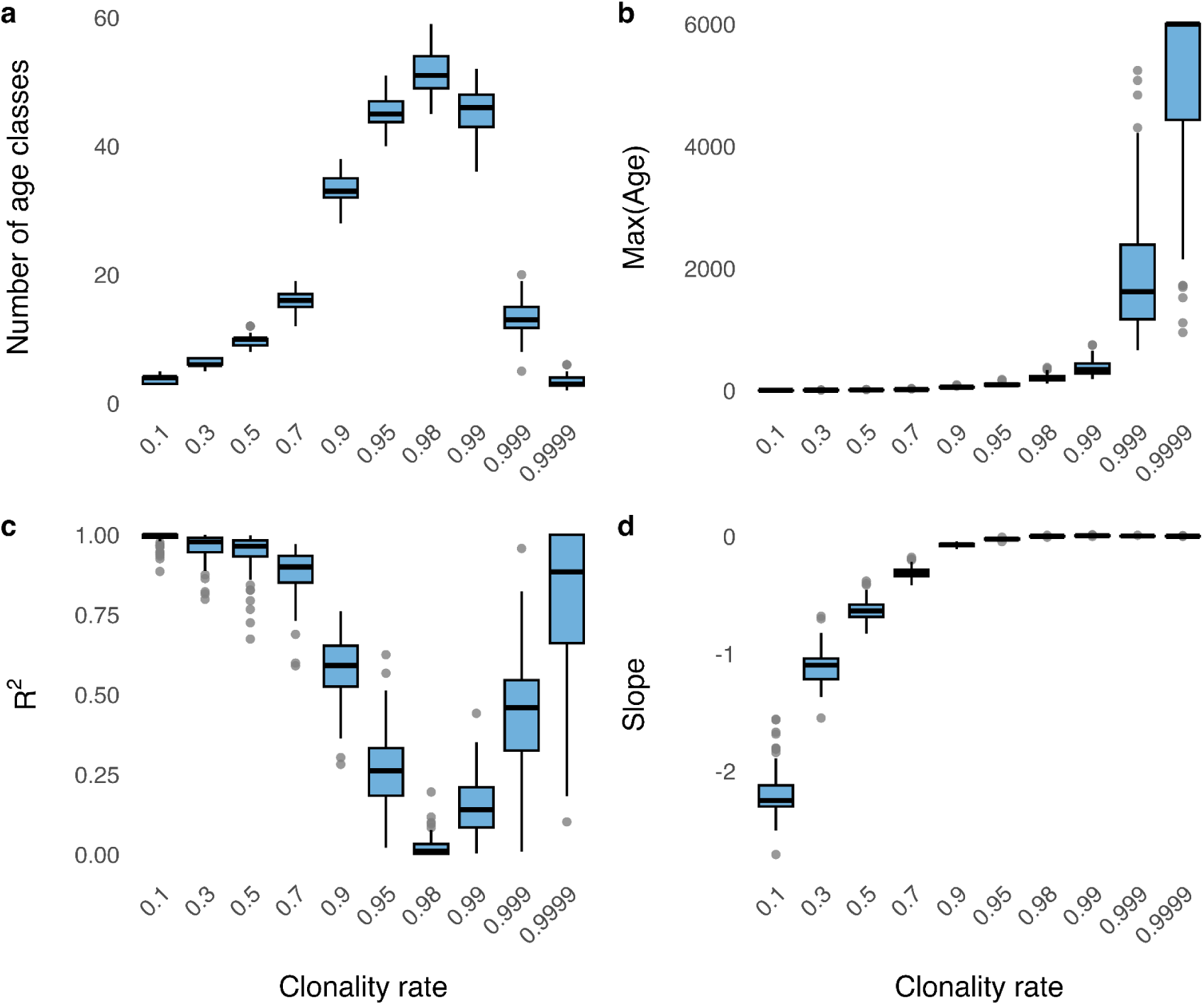
Clone age distribution metrics across the clonality rate gradient at generation 6,000. Boxplots show (a) number of age classes, (b) maximum clone age, (c) coefficient of determination (R²), and (d) slope; from regression analysis of the distribution of number of individuals and clonal ages across the clonality rate gradient. Each box represents results from 100 replicated simulations per clonality rate.

To delve deeper and to quantify the observed transition from structured to stochastic age/abundance patterns, we summarized the distribution of clone ages using four indices : (i) the slope and (ii) the coefficient of determination of the linear regression between the logarithm of clone abundance and clone age at generation 6,000, and (iii) the median, (iv) variance, and (v) the maximum age of clones. The last two indices highlight how clonality rates shape the longevity of clones, while R² and slope values capture the predictability and steepness of the age/abundance relationship (Fig. 6, S7).

The variance in clone ages dramatically increased at high clonality rates (*c* ≥ 0.999), indicating a wider range of clone ages, in line with the increase in maximum clone age that often reached the maximal value of 6,000 generations for *c*=0.9999. The coefficient of determination (R²) was consistently high at low clonality rates (*c* = 0.1 to 0.7), indicating well-predictable relationships. As the clonality rate increased beyond 0.7, R² gradually declined, becoming null at *c* = 0.98, indicating that the abundance of clone age was distributed randomly all over the possible clone age in different simulations. Interestingly, at the extreme clonality rates (*c* ≥ 0.999), the R² values increased again, implying a recovery of the age/abundance relationship. However, visual inspection of the scatter plots (Fig. 5) revealed considerable noise in the age/abundance relationship, the correlations being mostly driven by a few extremely old and abundant clones that were present in all simulation replicates (Fig. S4).

Similarly, the slope of the regression line was negative and steepest under low clonality rates, reflecting an exponential decrease in clone abundance with increasing clone age. This slope became flatter (less negative) as the clonality rate increased around *c* = 0.98, when the age and abundance became uncorrelated. At the higher clonality rates (*c* ≥ 0.999), the slopes are null. Together, these results highlight that the strength and sign of the age/abundance relationship vary considerably on either side of a tipping point of maximal unpredictability of clone age distribution.

### Summaries of clone age distribution correlate with some population genetic indices

We then examined how the distributions of abundance of clone age impacted population genetic indices, using Spearman correlations among metrics of clone age distributions (Fig. 6) and the value of population genetic indices (Fig. 3), both calculated at generation 6,000 and for simulation data obtained with μ = 10⁻^3^ (Fig. 7).

**Fig. 7.**
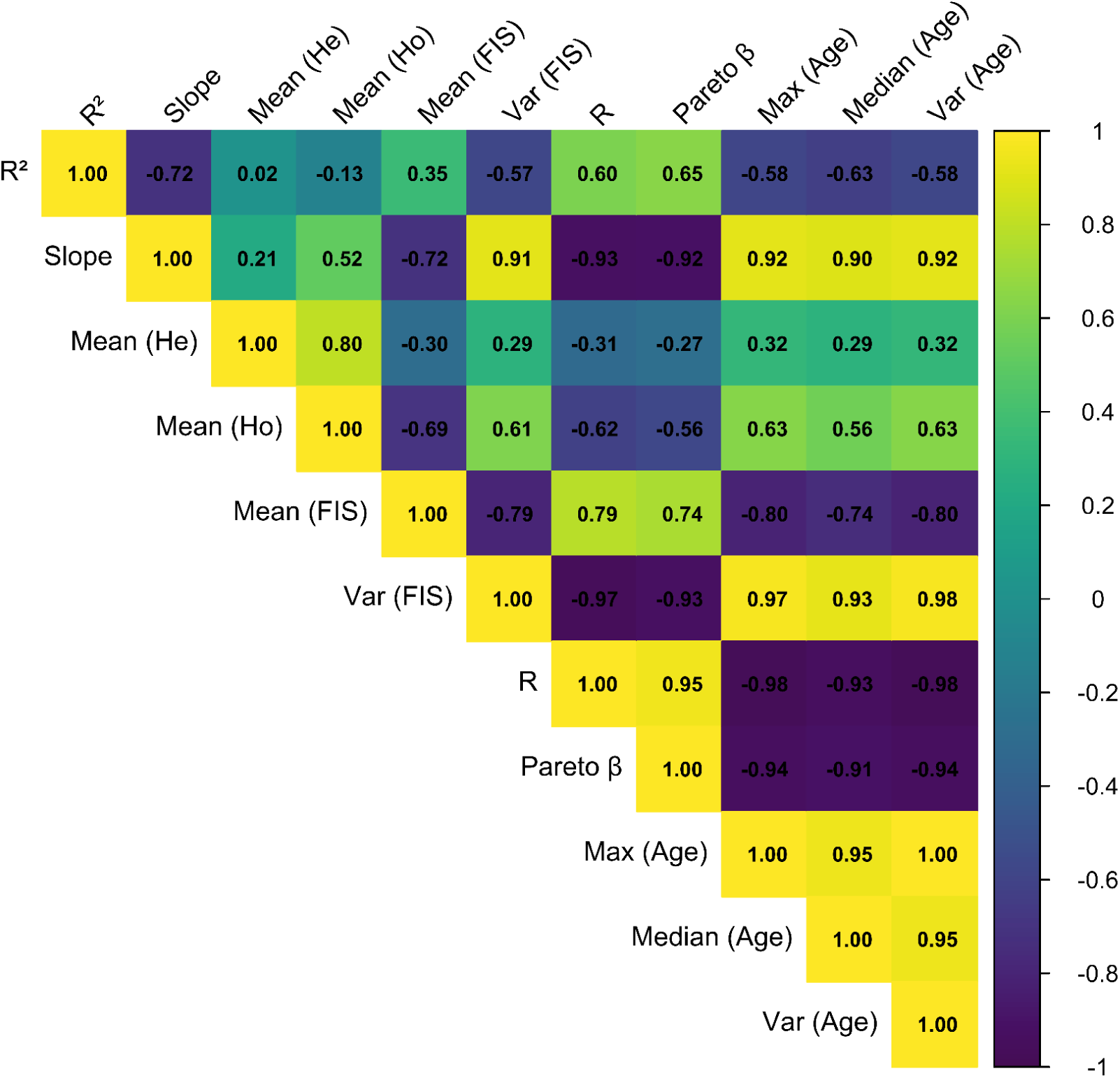
Relationship between clone age distribution metrics and population genetics indices at generation 6,000. Spearman correlation matrix among population indices and clone age distribution metrics at generation 6,000. Variables include: R² and slope (from clone age-size regressions), observed heterozygosity (*H*_O_), expected heterozygosity (*H*_E_), mean and variance of inbreeding coefficient (*F*_IS_), clonal richness (*R*), *Pareto β* (a measure of clone size inequality), and the maximum, median, and variance of clone ages. Each cell shows the correlation coefficient between two variables, color-coded from purple (–1) to yellow (+1).

Both R² and slope are correlated with clonal richness (R): R² correlates positively (ρ = 0.60) while slope correlates highly negatively (ρ = –0.93), and similarly with Pareto β (ρ = 0.65 and –0.92, respectively). As expected, the simulated populations with younger clones maintain higher genotypic diversity (high *R*) or have more even clone size distributions (high *Pareto β*). Spearman analysis also revealed a positive association (ρ ≥ 0.90) linking the three clone age longevity measures (maximum age, median age, and variance of age) to clonal richness (*R*) and to the *Pareto β*, indicating that populations containing very old clones also harbour fewer genotypes, but more evenly abundant. The slope of the age/abundance relationship exhibited a strong positive correlation with all longevity measures (ρ ≈ 0.92), consistent with rapid clone turnover limiting the build-up of ancient clones.

Classic population genetic indices were more weakly connected to the age structure: expected and observed heterozygosity showed modest correlation to clone age measures. Mean and variance of *F*_IS_ displayed a moderately high correlation with longevity metrics.

Mean *F*_IS_ shows weak positive correlation with R² (ρ = 0.35) and moderate negative correlation with slope (ρ = –0.72), whereas variance of *F*_IS_ is associated with both R² (ρ = –0.57) and slope (ρ = 0.91). Both heterozygosity indices (*H*_O_ and *H*_E_) display weak correlations with R², as well as moderate correlations with the regression slope.

Overall, the correlations show that longevity measures are most strongly linked with clonal structure (*R* and *Pareto β*) and the inbreeding index (*F*_IS_), indicating that the persistence of a few long-lasting clones in the population reduces the number of distinct clones and increases genetic similarity among individuals. In contrast, their correlations with genetic diversity among or within individuals (*H*_E_ and *H*_O_) are comparatively weak, highlighting that these metrics are less sensitive to clonal age structure.

### Examining clone age metrics clarifies variability in population indices

Then we investigate how clone age structure influences overall population genetic diversity, with an emphasis on the variability across simulation replicates. The relationships are illustrated for the slope and R^2^ of the clone age distribution (Fig. S5, 8) and for the maximum clone age (Fig. S6) with key genetic indices across clonality rates, at generation 6,000. As illustrated above, the variation in slope values correlates very well with genotypic diversity indices along the clonality gradient (Fig.8). Nevertheless, for low values of *c* (*c* ≤ 0.5), we can observe a lack of correlation among simulation replicates of the same modality. This is very clear for *c* = 0.1 or 0.2, for *Pareto β* (Fig.8f). The variance between replicates for Pareto B does not correlate with differences in slope values, which indicates that evenness of genotype abundance is not linked to differences in clone ages (Fig. S5).

**Fig. 8.**
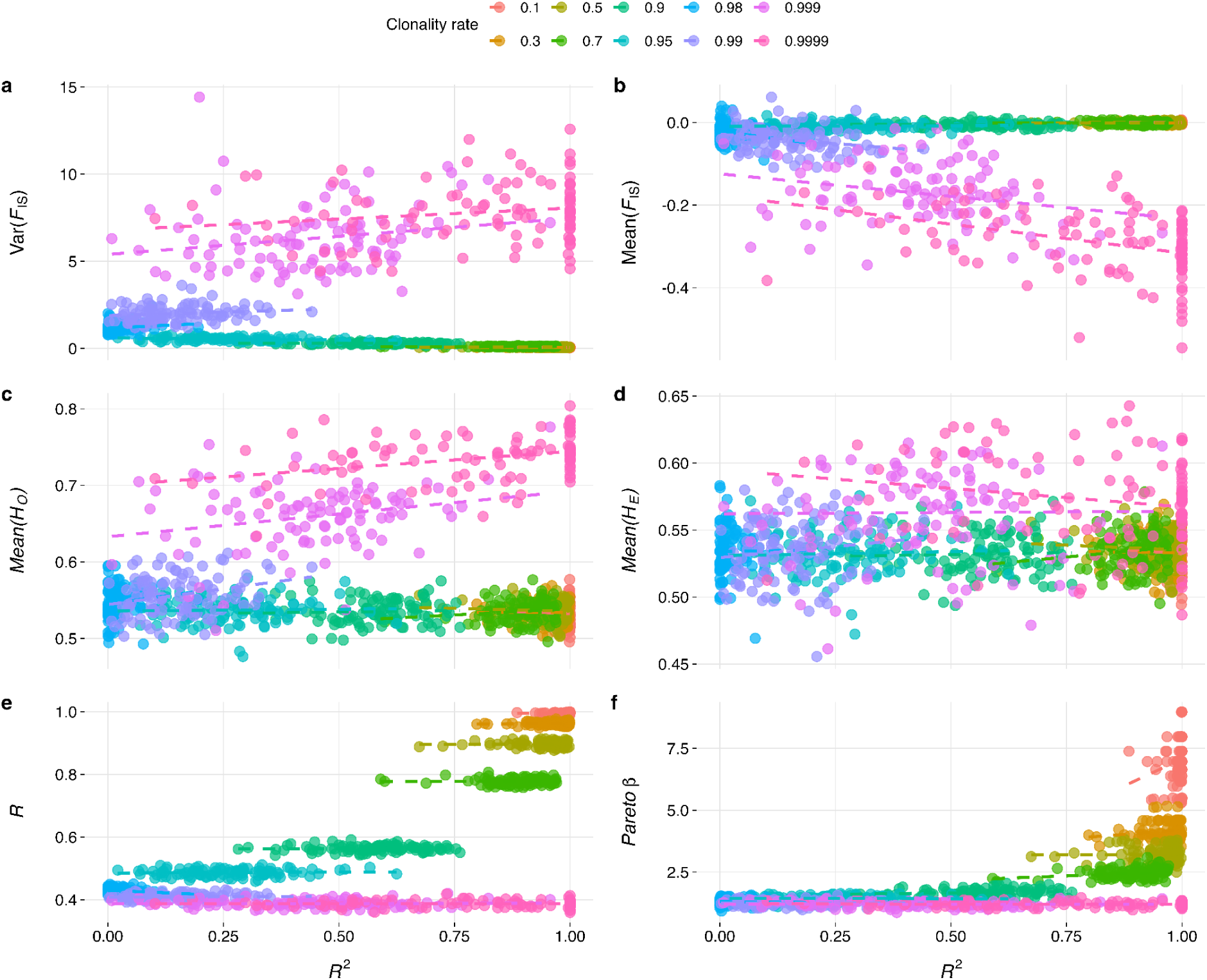
Relationship between clone age distribution R2 and population genetics indices across a gradient of clonality rates at generation 6,000. Scatterplots of the R^2^ of the semi-log regression between clone age and log-transformed abundance (x-axis) against six indices (y-axes) computed for each replicate and clonality rate: (a) variance in *F*_IS_, (b) mean *F*_IS_, (c) mean observed heterozygosity (*H*_O_), (d) mean expected heterozygosity (*H*_E_), (e) clonal richness (*R*), and (f) *Pareto β*. Each point represents one of 100 simulation replicates at a given clonality rate. Dashed lines show the best-fit linear regression for each clonality rate distribution.

The other metrics that summarize clone age distribution (R² of age/abundance relationship and maximum age) seemed to better explain the variation among simulation replicates, especially for genetic diversity and inbreeding coefficient for high rates of clonality. In particular, there are positive correlations between *H*_O_ and R² or maximum age for *c* ≥ 0.99, which indicates higher heterozygosity with increased clone age among replicates. In parallel, we observed negative correlations between mean *F*_IS_ and these metrics of clone age distribution. One final correlation worth noting is the one between var (*F*_IS_) and R², but var (*F*_IS_) is not correlated with maximum age. For this relationship, the scatter plots for *c* ≥ 0.99 are overlapping. Here, we clearly denote the effect of heterogeneity in clone age distribution among simulation replicates on this key parameter, which is associated with more or less rare recombination events, by chance.

## Discussion

Our theoretical analysis shows that patterns of clone age structure vary with the rate of clonality, and that different clone age distributions, in turn, influence population genetic indices. Our results help understand that variations in clone age distributions may explain part of the large ranges of population genetic values found in highly clonal populations. Altogether, these results shed new light on how to better infer clonal structure in natural populations.

### Identification of a tipping point between two processes ruling the birth and death of clones

The most original aspect of our analysis is the identification of a numerical tipping point in the change of clone age distribution. As the clonality rate increases, we observe a shift from a perfectly compact and determined age structure to completely variable and diffuse age distributions across replicates. From this tipping point onwards, we observe the emergence of old (surviving hundreds of generations), even very old (surviving thousands of generations), clones, which mix with younger ones. The age structure becomes less variable and spread as the proportion of old clones increases at high clonality rates, but clone age distributions remain highly variable from one simulation to another. This result illustrates, if need be, the high degree of stochasticity in highly clonal populations and the fallacy of adopting deterministically-based interpretations of their evolutionary trajectories (Reichel et al., 2016). This tipping point, at which we observe a completely variable and diffuse age distribution, actually corresponds to two processes balancing each other out. At low clonality rates, the age distribution is driven by the rapid turnover of clones, whose lifespan is determined to be exponentially decreasing. Indeed, at each generation, the proportion of new genotypes created by sexual reproduction is high, while the number of genotypes that can be maintained by clonality in the next generation is low. At high clonality rates, the number of surviving clones also decreases, but due to the random fluctuation in clone abundance over time (an analog of genetic drift, but at the scale of genotypes rather than alleles (Janko et al., 2008). This chance of survival applies both for old and young clones, whose establishment and persistence are impeded by the dominance of a few (more or less old) clones. Accordingly, the tipping point is where there was the highest number of classes of clone age. Mathematically speaking, this tipping point amounts to a diffusion between the two laws that determines the birth and death of clones (Ethier & Kurtz, 1986; Ewens, 2004). In the latter, we discuss how these two processes shape the variation of population genetic indices, hence offering the opportunity for a better disentangling of the evolutionary trajectories of partially clonal populations.

### Turnover of clones over time as a function of rates of clonality without selection and migration

Turnover of clones in populations has been advanced as one of the hypotheses to explain the long-term persistence of clonality in some species and a key process to understand if clonal reproduction is an “evolutionary dead-end” for species (Butlin et al., 1999; Jokela et al., 2009). New clones would allow populations to escape the selective clonal decay of lineages as they age (Paland et al., 2005) or as they face external selective pressures such as antagonist coevolutionary arms race between hosts and parasites (Jokela et al., 2009). Our results showed that at low rates of clonality, in a finite mutating population, the lifespan of clones was very short, from a couple to a few dozen generations before being replaced by new ones. Populations needed to reach very high rates of clonality (c>0.99) before observing the mean lifespan of clones exceeding 100 generations. This high clonal turnover was just fueled by random fluctuation and loss of clones over generations. In a finite mutating partially clonal population, as long as clonal rates are below 0.99, we therefore expect a rapid succession of clones over time without invoking external selective pressures driven by Red Queen mechanisms, migration, or exponentially-growing demography, as respectively proposed by Jokela et al. (2009) Jokela et al. (2009), Janko et al. (2008), and Angaji et al. (2021). Even in a strict clonal population, as long as diversity in clones is maintained, a dynamical succession of clones is expected, as observed in cancer tissues, for example (Selich et al., 2016; Gallaher et al., 2019). In partial clonality, the process is even strengthened: sexual reproduction provides continuously new clones that have to struggle against the low odds to maintain and increase in frequency. As Butlin et al. (1999), we call the community to monitor and study more spatio-temporal series of partially clonal populations in natural conditions to document this clonal turnover and further disentangle the neutral vs. selected hypothesis of clone decay (Janko, 2014).

### Evolutionary trajectory for low *c* values

At low clonality rates, below the tipping point, we observe a gradual establishment of clonal structure, with more individuals becoming clonemates as clonality increases. In accordance, we observe a gradual decrease in both clonal richness (*R*) and evenness (*Pareto β*). The best metric that then summarises clone age structure is the slope of the regression between clone abundance and age, which well correlates in turn with the variation of genotypic indices. Notably, the variation in population genetic indices only concerns indices describing genotypic diversity. At low clonality rates, allelic diversity and its apportionment within and between individuals are almost unchanged compared to pure sexual reproduction. The mean inbreeding coefficient (*F*_IS_) remains close to zero as the population follows Hardy-Weinberg expectations under near-random mating. Similarly, the variance in *F*_IS_ is increased compared to strictly sexual populations, but remains low, reflecting the homogenizing influence of sexual reproduction on the genetic structure of the population towards genotype frequencies expected under Hardy-Weinberg’s assumptions. One sexual reproduction event every 100 generations is enough to erase traces of clonal multiplication on allelic diversity and its apportionment within and between individuals (Rouger et al., 2016). At low clonality rates, the pace of clonal turnover imposes transient clonal structure, as a dynamical process dominated by sexual reproduction that continuously reshuffles alleles into new genotypes that may survive as clones for a few generations only, before being replaced by new ones.

### Evolutionary trajectory for high *c* values

Conversely, as clonality rate exceeds the tipping point (c ≥ 0.98), populations enter a different evolutionary dynamic characterized by increased unpredictability of clone age distributions that mirrors large variation in population genetic structure (Reichel et al., 2016; Stoeckel et al., 2021). The variation in genotypic indices (clonal diversity, *R,* and evenness, *Pareto β*) is no longer informative since they live very close to their minimum values. In addition, the variation in allele-based indices (*H*_O_, *H*_E_, *F*_IS_) lies in a range of values typical of high clonality with a very high overall excess of heterozygotes and a high variance in these indices across loci (Balloux et al., 2003; Halkett et al., 2005; Stoeckel & Masson, 2014). Differences in clone age distribution, due to a slow turnover of clones, making their aging and survival very stochastic, become the main driver of variability in genetic indices, which explains why populations with the same rate of clonality can have different genetic signatures. These stochastic trajectories pose major interpretation challenges for applied studies, as the range of possible values of population genetic indices increases and measured values in studied populations lose their bijective reliability at high rates of clonality (Stoeckel et al., 2021). Here, we show that part of this stochasticity, observed for high clonality, is explained by differences in age structure between simulation replicates. Notably, the mean *F*_IS_ correlates negatively with the maximum age of clones, and the variance of *F*_IS_ correlates negatively too with the coefficient of determination of the clone age/abundance relationship. These correlations result from different proportions of old and young clones in different populations, with older clones tending to increase individual heterozygosity (*H*_O_) and heterogeneity in genetic composition (variance of *F*_IS_) within the population. Beyond the tipping point, we clearly document the effect of clonal drift combined with a limited effect of mutation accumulation at µ =10^-3^ on the evolution of highly clonal populations.

### Distortion of evolutionary time for high *c* values and empirical consequences

As expected (Reichel et al., 2016; Stoeckel et al., 2021), the temporal change of classical population genetic indices shows that it takes much more time at high rates of clonality to reach steady-state values. This is especially true for the dynamics of *F*_IS_ and leads to the very large disparity between the evolutionary trajectories of low or highly clonal populations. As a result, it is difficult to compare the variation of population genetic indices, all else being equal (in time). Typically, the mutation-drift equilibrium depends on the value of *c* (Reichel et al., 2016), which results in a distortion in time along *c* values. This strong interplay between time (left so that clonal drift has acted), rate of clonality, and values of population genetic indices is difficult to grasp, but can be easily explained by the differences in turnover between clones, depending on the rate of clonality. Up to values of *c* = 0.99, the age of the oldest clones is negligible compared to the duration of a simulation run. However, this is no longer true at higher *c* values, with an increasing proportion of old clones, which can by chance reach the maximal age set to the duration of the simulation run. The disadvantage of such a long simulation duration is that it leaves room for genetic drift to fix alleles if mutation does not compensate for the loss of polymorphism. Typically, this leads to artifactual values of population genetics indices for very low mutation rates (*µ* = 10⁻⁶). Because of these extremely slow dynamics and these artifacts, it is difficult to compare theoretical expectations with empirically observed values for low mutation rates. Therefore, we advocate using highly variable molecular markers, such as microsatellite loci, when monitoring partially clonal populations sampled *in natura*.

### Insights for inferring clonality rates

Our results clearly underline the complexity of inferring clonality rates from population genetic indices. The inference of *c* is hindered by a double paradox regarding the variation of population genetic indices along clonality rates.

At low clonality rates, the variation in clonal richness R calculated on all the individuals of a population is fully determined by the *c* value (equation 8 in Stoeckel et al., 2021), which would naively suggest that the values of *c* can be easily identified. Unfortunately, the R index is highly biased and subject to strong variability when considering subsamples of the total population. Our study shows that, unfortunately, the age distribution is quite noisy too, at low *c* values, as evidenced by the variability of the slope of the clone age/abundance relationship. No index derived from the age distribution can be confidently used to infer low *c* values. Worse still, the age structure does not correspond to the relative abundance of genotypes captured by the Pareto B index, which adds to the complexity of analysing such transient clonal structure. Therefore, contrary to what might be expected in a deterministic case (Burt et al., 1996), we cannot use clone resampling to accurately estimate small values of *c* (Ali et al., 2016).

At high clonality rates, the issue of inferring *c* value is reversed. The index that correlates best with the variation in *c* is *F*_IS_. With little bias, this index is extremely robust to variation in sample size, but unfortunately, it shows high variability between simulations for the same values of *c*, as expected from the study of theoretical distributions obtained analytically (Stoeckel & Masson, 2014). The inference of *c* is hindered here by the high stochasticity of evolutionary trajectories in highly clonal populations. Our detailed analysis shows, however, that part of this variability can be explained by differences in the age structure of clones, particularly through differences in the relative abundance of young and old clones. This signal, linked to clonal drift, could provide additional information in combination with the signal provided by changes in allele frequencies through time, which is already used in the Bayesian method ClonEstiMate (Becheler et al., 2017).

Ultimately, our simulation study indicates that clone age structure is more insightful in deciphering long-lasting clonality rather than transient genotype turnover. This property can be used to better understand in more detail the origin of clonal structures in species with mixed reproduction, typically those many cases of coexistence of sexual and clonal lineages (Halkett et al., 2005; Abdalrahem et al., 2025). In this case, an important biological question is whether all individuals form a single population (as in our simulations) or whether there is additional reproductive isolation segregating sexual and clonal lineages (such as determinism for the loss of sex). Our study shows that the observation of persistent clones is indicative of a very high rate of clonality, even if these clones are rare within a sample. Typically, we have shown that the survival of clones over more than 100 generations is incompatible with clonality rates lower than c = 0.9 without selection. In case of rare long-lasting clonal lineage, however, the use of current clonality inference methods (Ali et al., 2016; Becheler et al., 2017) would have led to the inference of a low rate of clonality, considering wrongly that all sampled individuals form the same population. This type of error has serious consequences for (mis)understanding the ecological and evolutionary dynamics of partially clonal populations. On the contrary, reasoning by reductio ad absurdum would enable us to demonstrate that there is a partition within the population between sexual and clonal lineages. Obtaining temporal samples that are sufficiently spaced out over time can seem like a major challenge. However, advances in ancient DNA analysis (Der Sarkissian et al., 2015; Maixner et al., 2021), coupled with robust methodologies for identifying clones based on genetic data sets (Stenberg et al., 2003; Halkett et al., 2005; Arnaud-Haond et al., 2007) now make it possible to analyze ancient samples from museum collections or herbaria and contribute deciphering reproductive modes in addition to study past demography and biodiversity in response to environmental changes (Orlando & Cooper, 2014). Adapted to an adaptive evolution framework, this work could provide a better understanding of the dynamics of the emergence and replacement of clonal lineages, which are, for example, often associated with disease outbreaks.

### Broader implications for the study of the evolution of reproductive modes

More broadly, our work on clone age distributions, discussed here only from a neutralist perspective, may have important implications for studies scrutinizing the selection of reproductive modes. A consensus has formed that the evolution of sexual reproduction is not determined by a single factor, but is in fact based on several processes that interact to determine the evolutionary successes of sexual or clonal lineages (Neiman et al., 2017). Among these, a key aspect lies in the temporal fluctuations of frequency-dependent mechanisms that bring these lineages into interaction (e.g., Pierre et al., 2022). As with any co-evolutionary process, the outcome of interactions depends on the direct or indirect relationships ruling the negative frequency-dependent selection among partners (Brown & Tellier, 2011). Here, time is of considerable importance because it allows for bypassing an immediate response caused by direct frequency dependence. Like the phenomena of dormancy or seed banks, which stabilize coevolutionary cycles (Tellier & K. M. Brown, 2009), clonal reproduction allows past adaptive phenomena to be stored in memory and causes a time shift that buffers the response to selection. This response will, of course, depend on the turnover of clones, the properties of which we have examined here. The current limitation of models of sexual reproduction evolution is that they often consider a game between two pure strategies: strict sexuality or strict clonality. Considering partial clonality amounts to extending the analysis to mixed strategies, which often allow for reaching better evolutionary trade-offs (Gross, 1996; Goodwillie et al., 2005; Barrett, 2010). Here, our analyses show that the effect of the clonality rate is not linear, and the emergence of a stochastic pattern of clone ages could contribute to an adaptive response by mixing young and old clones, each responding to selective events of varying time. These analyses would provide a better understanding of the conditions for the evolutionary emergence of complex life cycles beyond the debate on the prevalence of sexual reproduction versus purely asexual reproduction.

## Data availability

Python codes for model simulations and summary statistics are accessible at https://www6.rennes.inrae.fr/igepp_eng/Productions/Software.

R scripts used to analyse simulated data are available on GitHub at: https://github.com/ammarabdalrahem/clone_age_distribution.

## Acknowledgements

We thank Scott Heslop for proofreading an earlier version of the manuscript. This research was supported by grants from the French National Research Agency (ANR-18-CE32-0001, Clonix2D project; ANR-23-CE20-0032, ENDURANCE project). Ammar Abdalrahem received support through a PhD fellowship from the French Ministry of Education and Research (MESR). We used DeepL, Grammarly, and Perplexity to proofread, suggest synonyms, check grammar and spelling, and ChatGPT to help with R scripts. All suggestions from these tools were carefully reviewed and verified by the authors.

## Author Contributions

AA, SS, and FH conceived and designed the study. SD, PF, and FH supervised AA’s thesis. AA and SS produced the code and ran the simulations. AA and FH analysed the data and prepared the manuscript. CN, as the personification of the academic community, contributed to developing the methodological framework and the state of research. All authors revised and approved the manuscript.

---

**Fig. S1.**
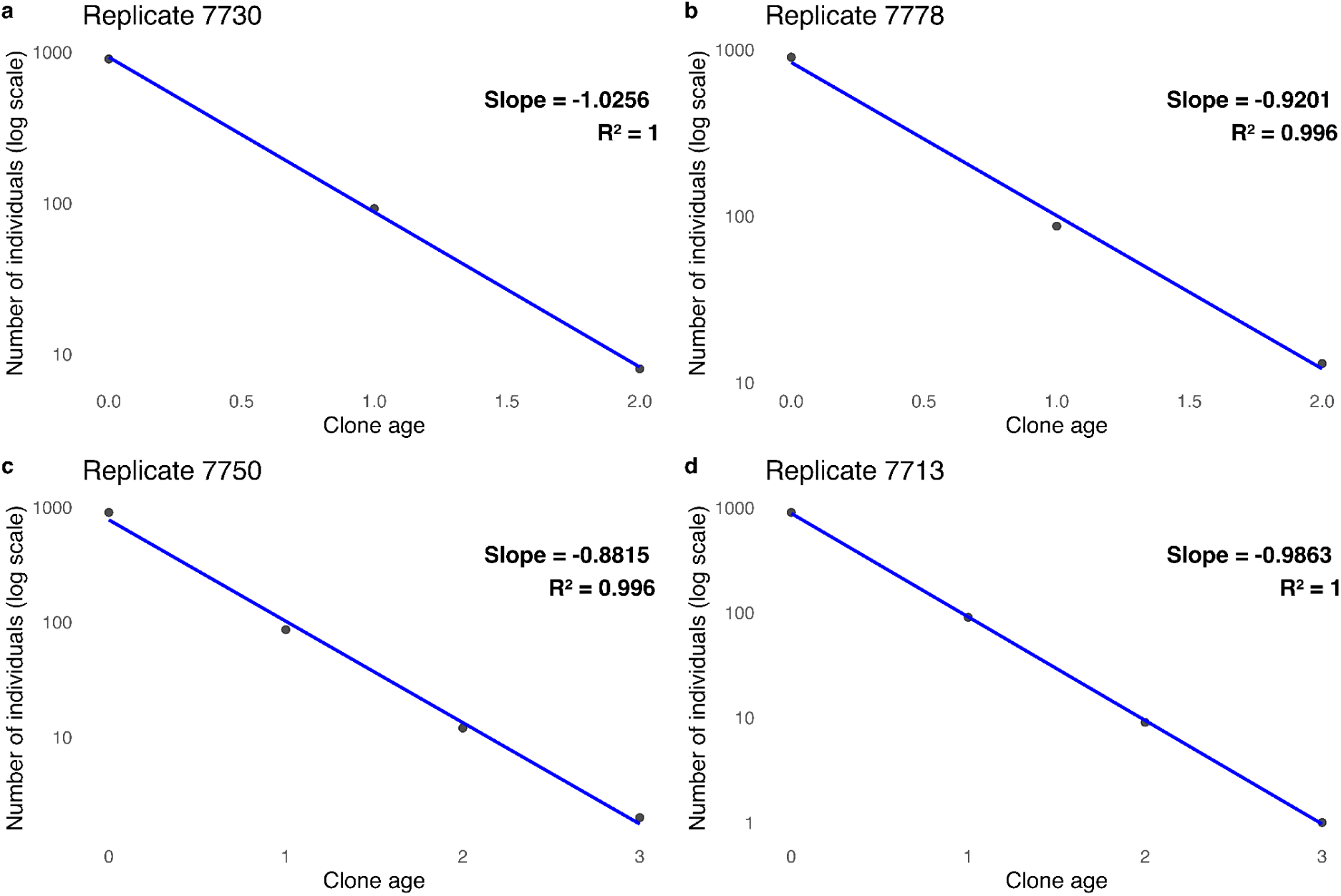
Distribution of clone ages at generation 6,000 with a clonality rate of 0.1. Scatterplots display the relationship between clone age and the number of individuals per clone (log scale) for four random simulation replicates with a clonality rate of 0.1, at generation 6,000. Each blue line is the best-fit regression, with annotated slope and R² values.

**Fig. S2.**
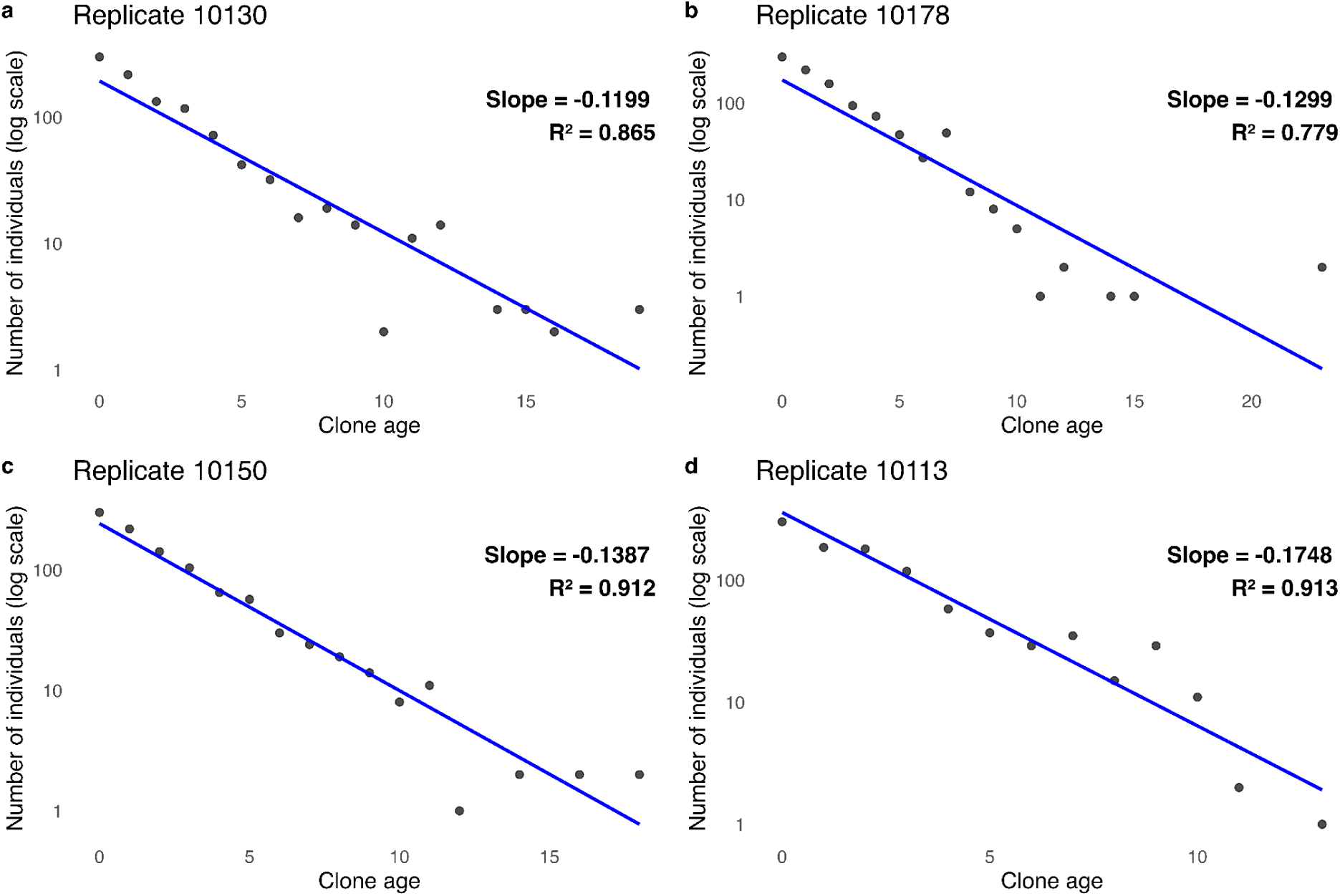
Distribution of clone ages at generation 6,000 with a clonality rate of 0.7. Scatterplots display the relationship between clone age and the number of individuals per clone (log scale) for four random simulation replicates with a clonal rate of 0.7, at generation 6,000. Each blue line is the best-fit regression, with annotated slope and R² values.

**Fig. S3.**
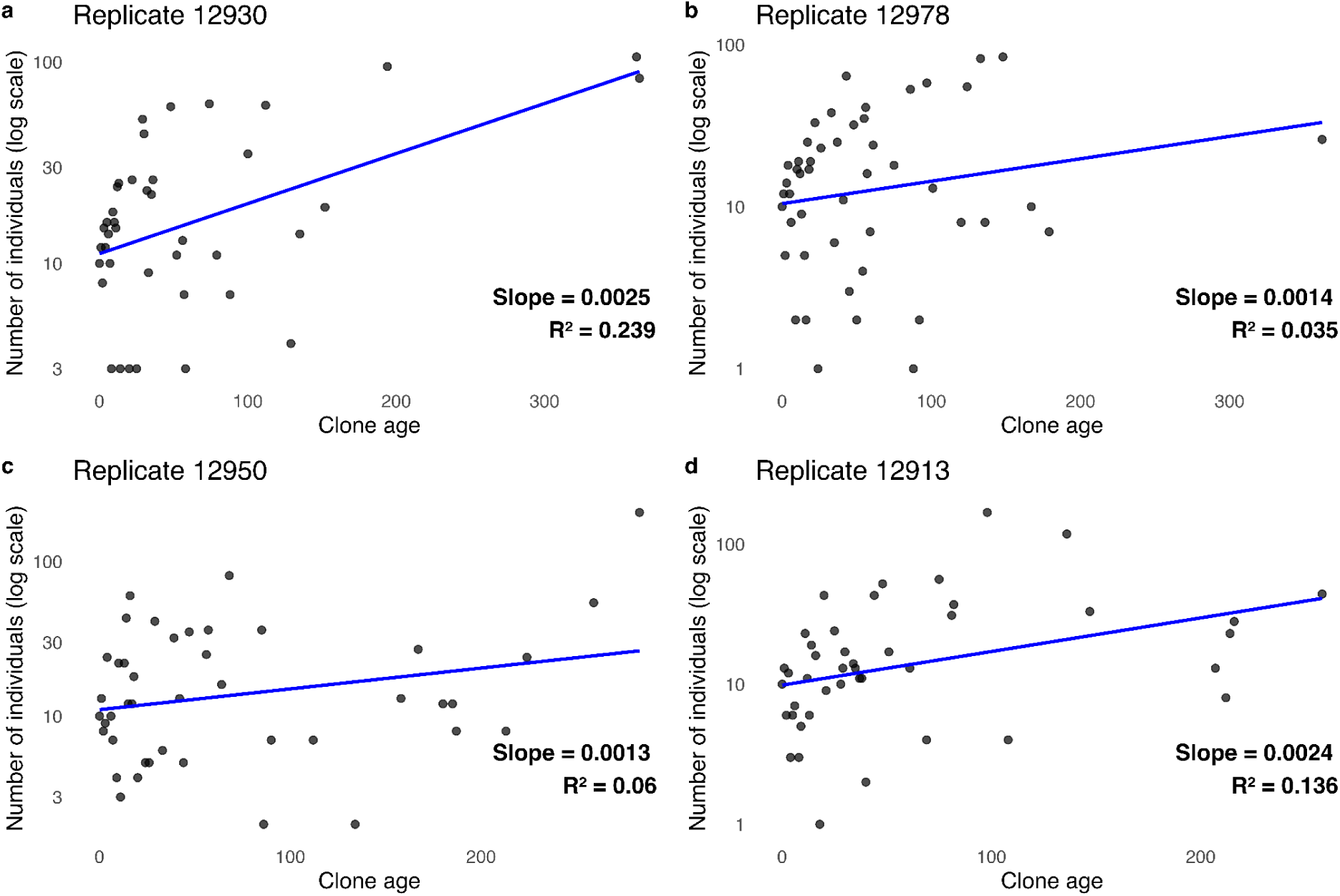
Distribution of clone ages at generation 6,000 with a clonality rate of 0.99. Scatterplots display the relationship between clone age and the number of individuals per clone (log scale) for four random simulation replicates with a clonal rate of 0.99, at generation 6,000. Each blue line is the best-fit regression, with annotated slope and R² values.

**Fig. S4.**
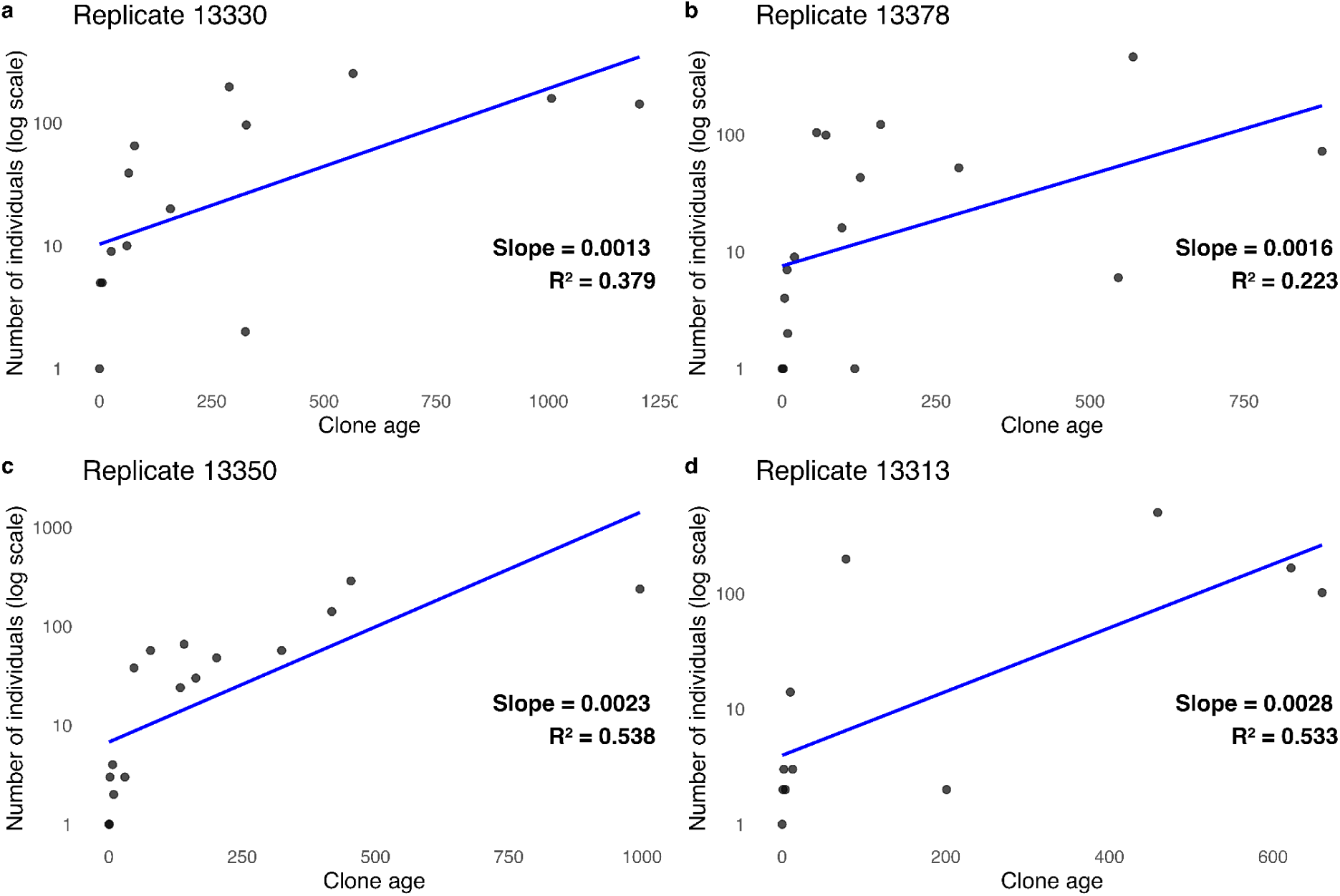
Distribution of clone ages at generation 6,000 with a clonality rate of 0.999. Scatterplots display the relationship between clone age and the number of individuals per clone (log scale) for four random simulation replicates with a clonal rate of 0.999, at generation 6,000. Each blue line is the best-fit regression, with annotated slope and R² values.

**Fig. S5.**
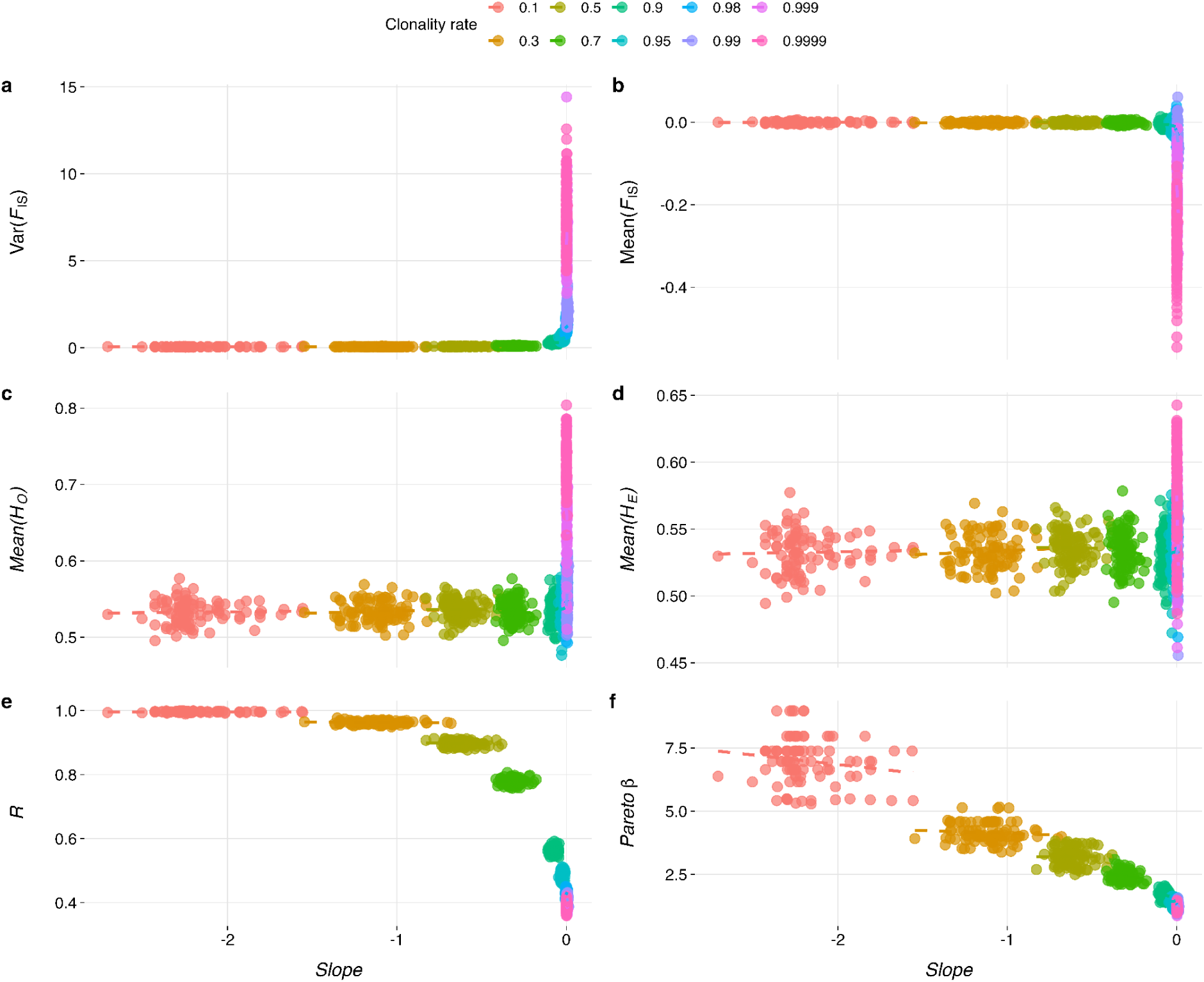
Relationship between clone age distribution slope and population genetics indices across a gradient of clonality rates at generation 6,000. Scatterplots of the slope of the semi-log regression between clone age and log-transformed clone size (x-axis) against six indices (y-axis) computed for each replicate and clonality rate: (a) variance in *F*_IS_, (b) mean *F*_IS_, (c) mean observed heterozygosity (*H*_O_), (d) mean expected heterozygosity (*H*_E_), (e) clonal richness (*R*), and (f) *Pareto β*. Each point represents one of 100 simulation replicates at a given clonality rate. Dashed lines show the best-fit linear regression for each clonality rate distribution.

**Fig. S6.**
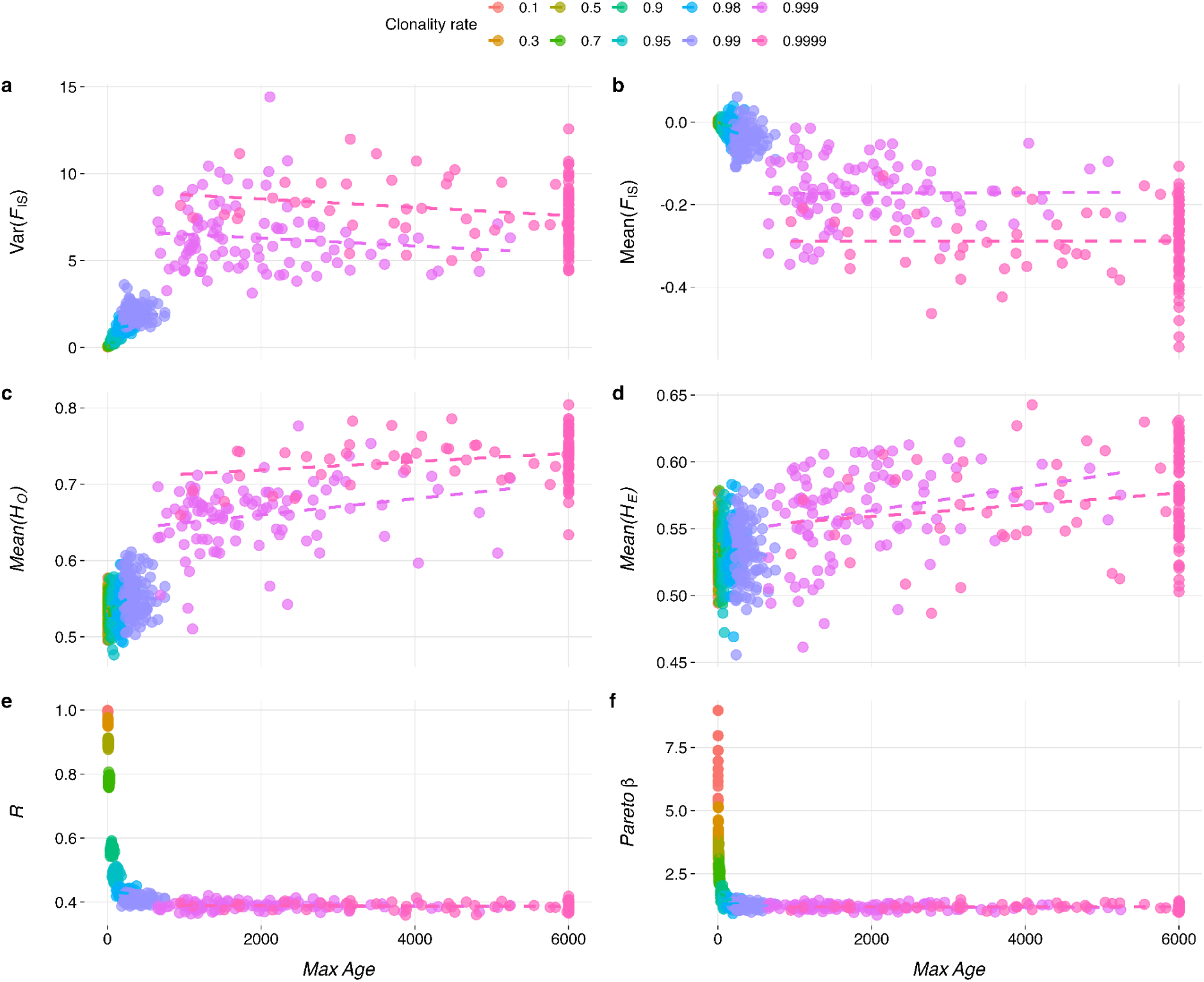
Relationship between the maximum of clone age distribution and population genetics indices across a gradient of clonality rates at generation 6,000. Scatterplots of the maximum clone age distribution against six indices (y-axes) computed for each replicate and clonality rate: (a) variance in *F*_IS_, (b) mean *F*_IS_, (c) mean observed heterozygosity (*H*_O_), (d) mean expected heterozygosity (*H*_E_), (e) clonal richness (*R*), and (f) *Pareto β*. Each point represents one of 100 simulation replicates at a given clonal rate. Dashed lines show the best-fit linear regression for each clonality rate distribution.

**Fig. S7.**
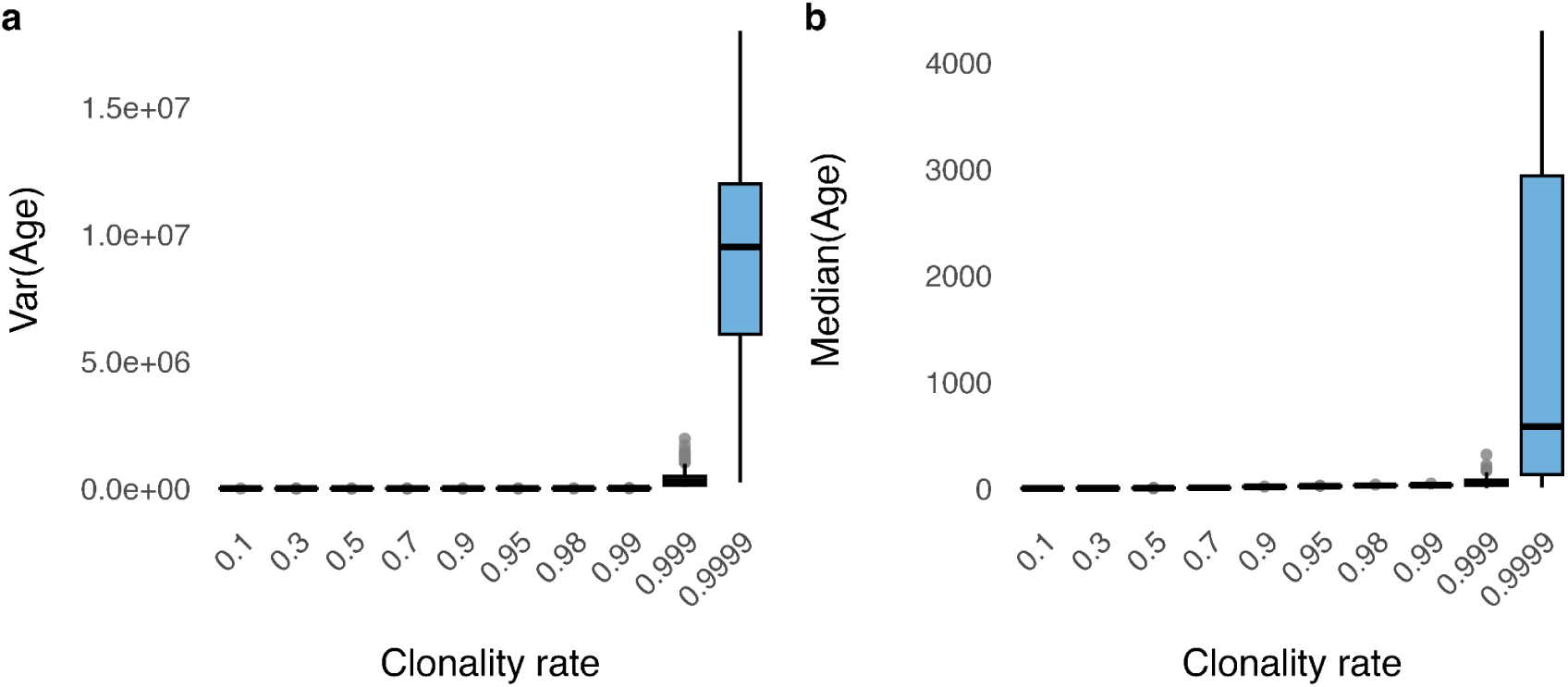
The variance and median of the clone age distribution across the clonality rate gradient at generation 6,000. Boxplots show (a) variance clone age, (b) median clone age. Each box represents results from 100 replicated simulations per clonality rate.

